# Quantity and quality of suitable matrices matter in reducing the negative effect of fragmentation on populations extinction risk

**DOI:** 10.1101/2020.08.10.244178

**Authors:** Bruno Travassos-Britto, Camila Hohlenwerger, José Miranda, Pedro Luís Bernardo da Rocha

**Affiliations:** Laboratory of Theoretical Ecology, Department of Ecology, University of São Paulo, São Paulo City, SP, Brazil; Laboratory of Landscape ecology and Conservation, Department of Ecology, University of São Paulo, São Paulo city; Laboratory of Biosystems, Institute of Physics, Federal University of Bahia, Salvador, BA, Brazil; Laboratory of basic and applied ecology, Institute of biology, Federal University of Bahia, Salvador, BA, Brazil

**Keywords:** Individual-based model, fragmentation, matrix use, population dynamics, survival time, extinction time, matrix heterogeneity

## Abstract

The negative effect of fragmentation is one of the main concerns in the study of biodiversity loss in landscape ecology. The use of the matrix has been considered an important factor because it can change the relationship of a population with the configuration of the landscape. A systematic way to assess the effect of matrix quality in fragmented landscapes could lead to a better understanding of how matrices can be used to suppress the negative effect of fragmentation. We built a computational individual-based model capable of simulating bi-dimensional landscapes with three types of land cover (habitat, suitable matrix and hostile matrix) and individuals that inhabit those landscapes. We explored in which situations changes in the proportion of the suitable matrix in the landscape and the degree of usability of this suitable matrix can mitigate the negative effect of fragmentation per se. We observed that (i) an increase in the matrix quality (increases in the suitable matrix proportion and/or usability) can suppress the fragmentation effect in 47% of the simulated scenarios; (ii) the less usable the matrix is the more of it is needed to suppress the fragmentation effect; (iii) there is a level of usability below which increasing the suitable matrix proportion does cause the fragmentation effect to cease. These results point toward a landscape management that considers the similarity of the matrix to the native habitat under management. We suggest that an index to measure the usability of elements of the matrix could be an important tool to further the use of computational models in landscape management.

## Introduction

Habitat conversion into human-dominated landscapes is one of the main causes of biodiversity loss worldwide (Maxwell et al 2016). Most of the negative effects of habitat conversion on biodiversity are known to be outcomes of the process of habitat loss and the process fragmentation on the species populations (Fahrig 2003, 2017, 2020. Fahrig et al 2019; Püttker et al 2020; Watling et al 2020). The negative effect of habitat loss is related to the reduction of the living area of species, which by its turn limits the size of populations living within the habitat, and smaller populations have a higher chance of extinction leading to increased loss of biodiversity and associated natural processes (Saunders et al. 1991). The negative effect fragmentation per se is explained by the fact that one large patch of habitat can maintain a larger population which has a small chance of extinction, while several small isolated patches of habitat can maintain only (if at all) very small populations which have a higher chance of extinction (Fletcher et al. 2018). As a consequence, fragmented landscapes support less populations than non-fragmented ones, even for the same amount of habitat.

Although there has been a lot of progress on the disentangling of the effects of habitat loss and fragmentation on biodiversity, many studies have been conducted under the notion of binary landscapes composed of a highly usable part (the habitat) and a highly hostile one (the matrix) (Fahrig, 1998; Saunders et al, 1991). However, it is well-known that human-dominated landscapes are complex environments encompassing habitat patches usually embedded in different types of matrices that may offer different degrees of usability to the species in the landscape (Boesing et al 2018; Chetcuti et al. 2021). This then creates a complex mosaic of land covers that goes beyond a binary landscape representation, including not only habitat patches, but also different matrix patches that may vary from completely hostile to very suitable for habitat species. Therefore, one important aspect of this discussion is understanding how the fragmentation effect per se is modulated by different aspects of the matrix (Ricketts, 2001; Ewers and Didham, 2006; Chetcuti et al. 2021). Considering the urgent call to reduce loss of species and processes due to human activities, understanding the role of the matrix is especially relevant for scenarios in which fragmentation has a negative effect on biodiversity.

Within human-dominated landscapes, two of the ways the matrix can favour the permanence of the population in the landscape are by offering complementary resources to the species and by increasing connectivity in the landscape (Ewers and Didham, 2006; Prevedello and Vieira, 2010). The matrix can provide resources that are no longer available within habitat fragments either because it is no longer there or because individuals are precluded from accessing it due to, for example, competition (Blitzer et al 2012; Driscoll et al 2013). In this case, even though the resource may not be of the same quality as that found in the habitat, it might be enough to increase the population’s abundance, thus favouring the population’s survival in fragmented landscapes. Moreover, patches of a more usable matrix can act as stepping-stones connecting fragments of native habitat that otherwise would be beyond the dispersal ability of individuals, even if these patches in the matrix cannot provide an essential resource by themselves (Ricketts, 2001; Rocha et al 2021).

However, different types of matrices may have different degrees of usability to habitat species, which then may shape the overall matrix quality for the community of species in the habitat (Prevedello and Vieira, 2010; Barros et al 2019). For example, Boeing et al. 2018 has shown that coffee plantations are more similar to Atlantic rainforests than eucalypt plantations or pasture and, therefore, offer less resistance to species movement and act as more suitable environments for rainforest species. Finally, the degree of usability of the matrix may also define the proportion of it that is needed in the landscape to ensure the population’s survival and thus mitigate the negative effects of fragmentation (Prevedello and Viera, 2010). For example, it is expected that low-quality matrices, compared to high-quality ones, may offer so little resource that an impressive amount of it may be needed to ensure the population’s survival in a fragmented landscape.

In this study, we move forward on the understanding of the effects of fragmentation per se by investigating the possible effects that matrices with different qualities may have on the population’s survival across fragmented landscapes. We used an individual-based model (IBM) to investigate how variations in the coverage and usability of a more suitable matrix can mitigate the negative effects of fragmentation per se on the population’s survival in the landscape.

## Methods

We created an IBM to investigate the scenarios in which the matrix quality can mitigate the fragmentation effect on the survival time of populations. The model can simulate a population of individuals of an ideal organism that can move, reproduce, and die with different probabilities depending on which type of land cover it is on. In this model, individuals inhabit a bi-dimensional landscape with three different types of land cover: habitat, suitable matrix and hostile matrix. We conceived the habitat cover type as an ideal environment where the focal organisms evolved to find the optimum set of elements that maximise their survival and reproduction chance. The hostile matrix was conceived as an environment in which organisms are highly exposed to death threats, have very little resource access, and have no means to establish a reproductive site. The suitable matrix was conceived as an environment where organisms can find some elements to support life and reproduction but not of the same quality or as frequently as they would find in the habitat environment.

A landscape can be generated with different levels of fragmentation and general matrix quality. Landscapes with different matrix qualities were generated with changes in the ratio between suitable and hostile matrices and/or with different usability levels of the suitable matrix. We analysed how landscapes with different values of fragmentation and matrix quality affect the survival time of populations of such individuals.

## Details of the model

The model we built to conduct our analysis is largely based on Fahrig’s (1998), which was used to demonstrate the theoretical effect of fragmentation per se. In her model, Fahrig creates a model capable of simulating a binary landscape of habitat and matrix, and individuals that can move, reproduce and die. She uses the model to simulate landscapes with different levels of fragmentation of the habitat type and analyses the impact of this fragmentation on population survival time.

Despite being a 20+ years old model, it is still relevant to explore the effects of fragmentation per se in an elegant and objective way. Furthermore, Fahrig’s model was used as the basis for a vast discussion about the effects of fragmentation and habitat loss (e.g. Fahrig, 2003; Villard and Metzger, 2014; Fletcher et al. 2018) and using her model as a basis, makes our results easier to compare to other results in this discussion. Therefore, before conducting our own analysis, we first checked if we obtained the same results as described in Fahrig (1998) when setting the models to the same parameters (see recovery results in Supporting information S1).

Our model is identical to Fahrig’s model except that in our model the matrix is conceived as a heterogeneous environment composed of two types of land cover (hostile matrix or suitable matrix). In this section we describe implementational aspects of the model we built following the ODDprotocol (overview, design concept, and details) for describing individual-based models (Grimm et al., 2010).

### Entities, state variables and scale

Our model space and time are measured discreetly. Entities were conceived within a hierarchy of three classes as follows: the lowest class of the hierarchy is the “individual”; the intermediate class is the “cell”, and the superior class is the “landscape”. The hierarchy is defined by spatial envelopment. Units pertaining to the individual class dwell in units of the class cell, and cells are the particles of the landscape class. Each of these classes has its own set of state variables (Table 1).

**Table 1.** List of parameters and respective values used in the model analysis for this study.

| Class | Parameter | Description | Value |
| --- | --- | --- | --- |
| Landscape | AREA | Number of cell units in the landscape | 900 (30x30) |
|  | COVER | Proportion of the landscape covered by habitat | 0.10 |
|  | FRAG | Fragmentation index of the habitat and suitable matrix | 0.01-1.00 |
|  | USABILITY | Usability of the suitable matrix | 10%-90% |
|  | N USABILITY | Usability level of the habitat | 100% |
|  | *COVERM* | Ratio between suitable matrix and total matrix area in the landscape | 0.00-1.00 |
|  | *NIND* | Starting number of individuals in the simulation | 500 |
|  | *H USABILITY* | Usability level of the hostile matrix | 0% |
| *Cell* | *cellType* | Type of landscape cover | habitat/<br>suitable matrix/<br>hostile matrix |
|  | *cellX* | Horizontal coordinate of the cell in the cell grid | 0-29 |
|  | *cellY* | Vertical coordinate of the cell in the cell grid | 0-29 |
|  | *maxOcc* | Maximum number of individuals in the cell per time-step | 10 |
| *Individual* | *Use* | Perception of the usability of inhabited cell | 0%-100% |
|  | *indX* | Horizontal coordinate of the individual in the cell grid | 0-29 |
|  | *indY* | Vertical coordinate of the individual in the cell grid | 0-29 |
|  | *maxDist* | Maximum distance that can be walked in a single time-step | 4 |
|  | *Mort* | Probability of death per time-step | 0.05-0.50 |
|  | *Move* | Probability of movement per time step | 0.50-1.00 |
|  | *rProb* | Probability of reproduction per time-step | 0.00-0.50 |

#### Landscape class

Aunit of this class is a bi-dimensional square object composed of a cell grid of AREAnumber of cells. A landscape always has a given number of habitat cells and suitable matrix cells determined by *COVER*and *COVERM* respectively. Cells that are from neither type are assigned as hostile matrix. Each landscape has a given level of aggregation between cells of the same type (*FRAG*). The landscapes have periodic boundary conditions (the righ and left, and top and bottom edges are joined continuously).

#### Cell class

Aunit of this class is a square object that can be occupied by up to maxOcc individuals at the same time-step. Each cell has the information of which position it is in the cell grid (coordinates *cellX* and *cellY*). In our model, each cell can be of one of the three different types of land cover: habitat, suitable matrix and hostile matrix.

#### Individual class

Individuals were conceived exactly like the ones described in Fahrig (1998). In each time-step, individuals interact with the cell in which they are. Each individual has a probability to die (*mort*), reproduce (*rProb*) and move to another cell (*move*). These three probabilities change linearly depending on the usability level of the inhabited cell (Figure 1). Higher usability results in lower probabilities of death and movement and higher probabilities of reproduction. The perception of the individuals is associated with the perception of the usability level of the cell they inhabit in the present (*USABILITY*). Each individual has parameters defining which cell they inhabit (coordinates in the cell grid *indX* and *indY*). An individual might move in a given time-step, and if it does, it will move a random distance up to maxDist cells in that time-step at a random direction.

**Figure 1.**
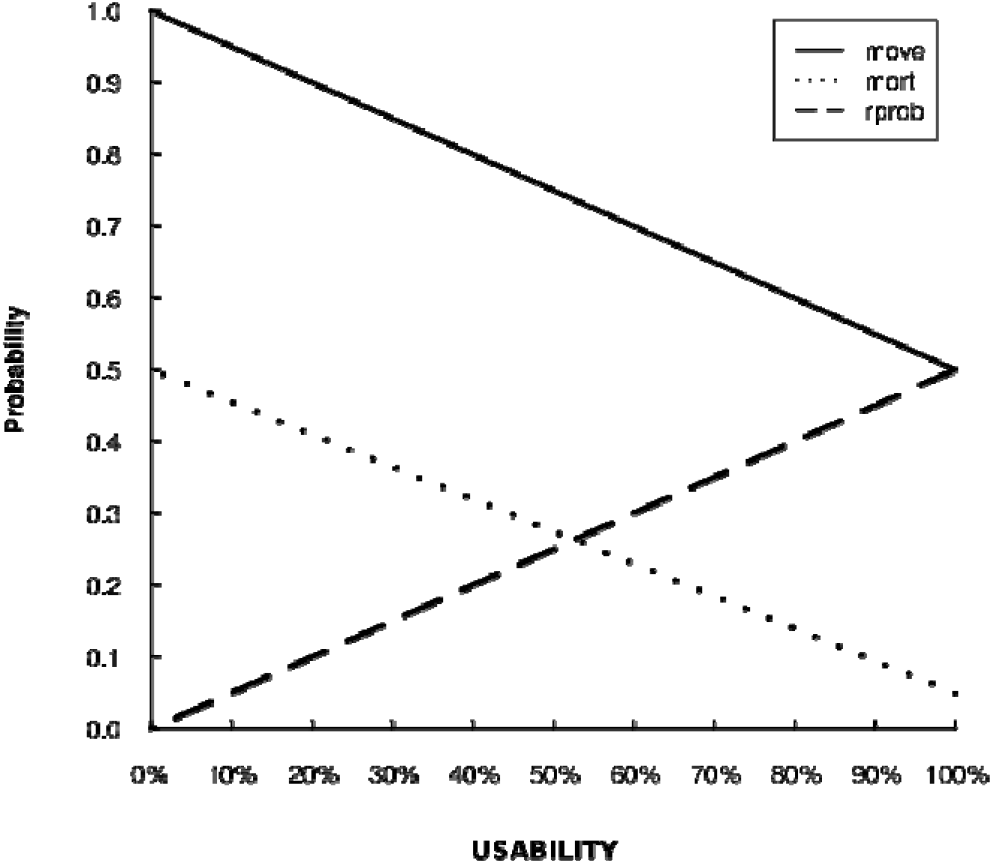
Relationship between the usability level of the cells and the probability of execution of movement (*move*), death (*mort*) and reproduction (*rprob*) by a single individual in one time-step.

### Routines

#### Landscape construction routine

Landscapes are defined by combinations of the following parameter values: *COVER, COVERM, USABILITY*, and *FRAGFRAG*defines the aggregation level of the habitat type cells and of the suitable type cells. We recreated the same algorithm used by Fahrig to implement fragmentation of breeding habitat in her model (see Fahrig, 1998). The difference in our model is that the algorithm applies to both habitat and suitable matrix. Primarily, all cells are defined as hostile matrix, and they will be assigned as habitat or suitable matrix in the course of this routine. The focal environment type is also selected randomly (habitat or suitable matrix). The cells are selected one by one and assigned with the focal land cover type. However, this assignment is conditioned to a single rule that might be ignored. The rule is that a selected cell is only assigned with the focal land cover type if at least one of the neighbouring cells is already assigned with that type. This rule induces aggregation of the cells of the same cover type. FRAGis the probability to ignore this rule, and assign the selected cell as the focal cover type independently of its neighbours. If FRAGequals 1.0, the aggregation inducing rule is completely ignored, and the cells will be assigned randomly as they are selected. However, if *FRAG*is too low, the rule is applied most times; thus, generating a pattern in which cells of the same types occur aggregated. If a cell is not assigned in the routine, its type is not changed, and another random cell is selected. If the selected cell is assigned, then the focal type of land cover changes, and the routine is repeated. This routine is repeated while the number of cells intended for both types of cover (defined by *COVER*for habitat and *COVERM* for suitable matrix) is not reached. In the end of the process the landscape profile will have a proportion *COVER*of habitat, and a proportion *COVERM* of suitable matrix and it will be ready to run its first time-step.

#### Time-step routine

In the beginning of each time-step, the cells are selected randomly one by one, and each selected cell executes its internal routine. In each cell, the same process is applied to individuals a follows: the individuals are selected randomly one by one, and each executes its own individual routine. When an individual is selected, it defines the order in which it will execute its three possible actions (die, move and reproduce). Due to the probabilistic nature of these events, one individual might die, move and/or reproduce in one time-step, but it also could do nothing once it is selected. After all individuals have been selected and have had the chance to execute their three actions, another cell is selected. After all cells have been selected, the outcomes of individual actions are updated as follows: individuals who have moved are placed in the new cells; individuals who died are deleted; and cells in which individuals reproduced receive one new individual for each successful reproduction. If there are any cells with more than *maxOcc* individuals, the individuals in these cells are randomly deleted until there are only *maxOcc* individuals. This event is the only implementation of a density-dependent process in the model. After this step, another time-step takes place. This routine is repeated until the number of individuals equals zero or the number of time-steps equals 500. The number of time-steps, *FRAG, COVERM* and *USABILITY* are then registered.

All routines and subroutines are described in detail, graphically in Supporting information S2.

### Analysis

Before we explain how we used the model to analyse in which situations the matrix quality can mitigate the fragmentation effect, it is essential that we clarify some terms used in the explanation.

“Landscape profile” is the set of parameters used to construct a given landscape unit. “Landscape configuration” is the specific picture of the landscape considering the relative positions of cells of each land cover type. As our model has probabilistic components in the construction of the landscape, the same landscape profile could generate landscapes with different configurations (see Figure 2). “Matrix profile” is the combination of the two parameters related to the amount of suitable matrix cover on the landscape (*COVERM*) and the usability level of these types of cover (*USABILITY*).

**Figure 2.**
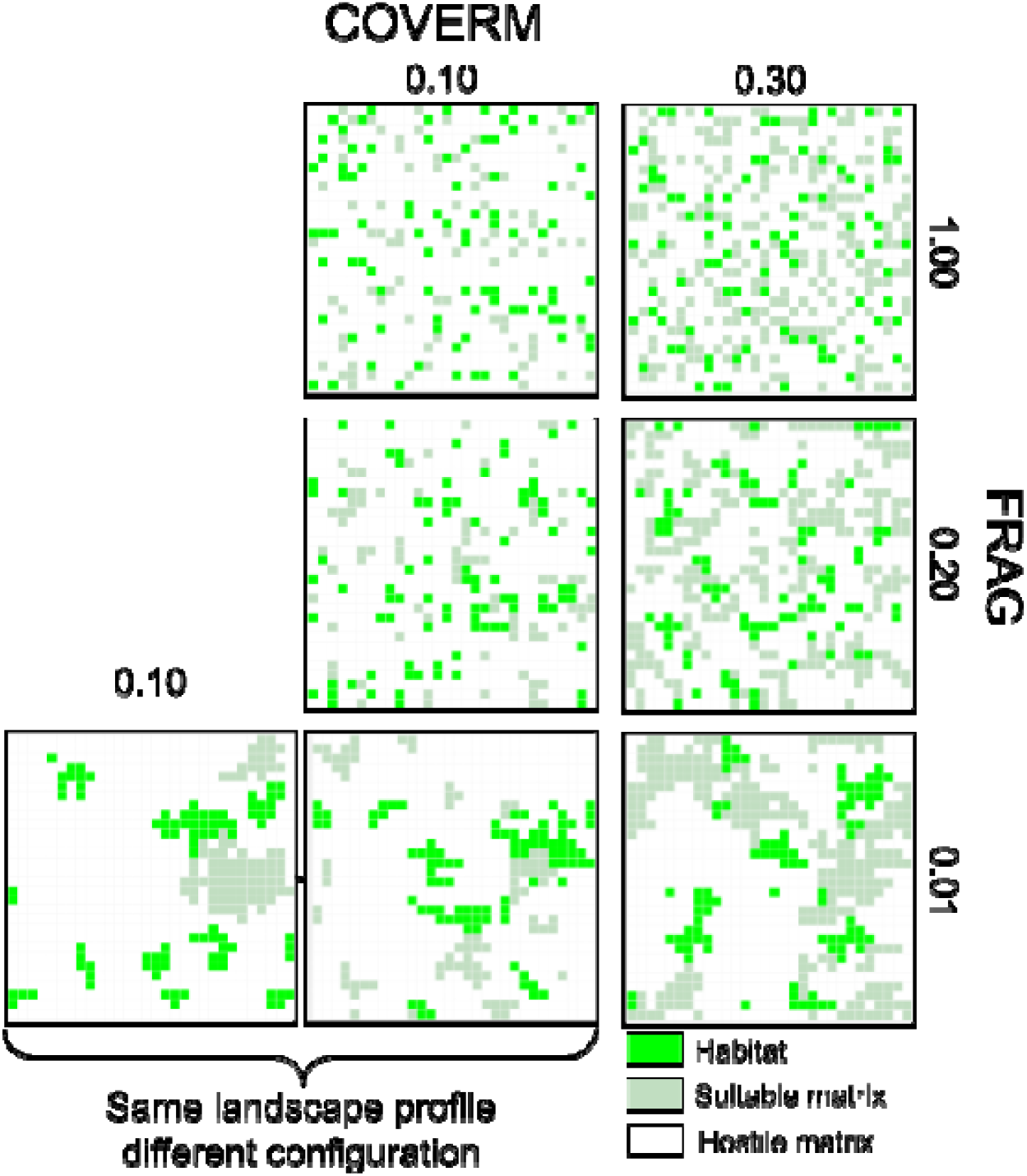
Graphical representation of landscapes generated with different levels of fragmentation (*FRAG*) and suitable matrix/total matrix area ratio (*COVERM*). These landscapes were generated using the same number of cells (*AREA*= 900) and the same proportion of habitat (*COVER*= 0.10). The landscapes united by braces were generated using the same parameters, but they present different configurations due to the probabilistic aspects of the landscape generation routine.

“Simulation” is a complete routine of creating a landscape with its set of parameters and individuals, and letting the individuals interact with the landscape until the population goes extinct or the number of time-steps reaches 500. In every simulation, the landscape profile is registered as the predicting variable and the time-step in which the population became extinct (or not if the simulation reached 500 time-steps) is registered as the dependent variable.

“Fragmentation analysis” describes the relationship between the level of fragmentation of the landscape and the survival time of the focal population. This analysis is conducted by running simulations at different values of *FRAG*and measuring how this changes the survival time of the population. The analysis conducted by Fahrig (1998) did not reveal any positive effect of fragmentation and, because our model is based on hers, we assume that the effect of fragmentation is only negative for this analysis. Therefore, the effect of fragmentation exists when for low values of *FRAG*the ideal population survives indefinitely (more than 500 time-steps) and for high values of *FRAG*, the population becomes extinct very quickly once the simulation is initiated resulting in a negative relationship (Figure 3). The fragmentation effect does not exist when one of two events occur: i) populations survive indefinitely independently of the level of fragmentation of the landscape; ii) populations become extinct very quickly independently of the level of fragmentation. In either case, the slope of the relationship between the fragmentation parameter and survival time would be close to zero.

**Figure 3.**
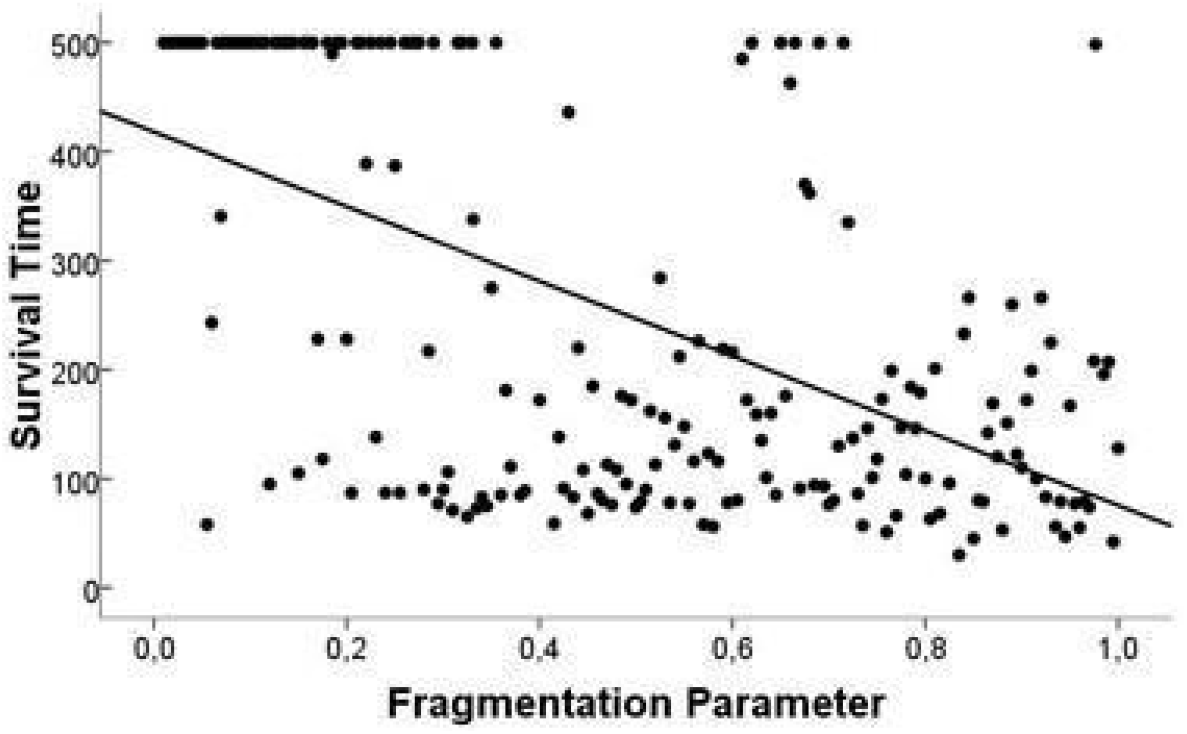
The negative fragmentation effect. This is the relationship between fragmentation and survival time when the entire matrix is covered by the hostile matrix type and all other parameters are held at default value. This graph is the result of 100 simulations, with each simulation run in a different value of the fragmentation parameter (*FRAG*= 0.01 -1.00).

A simulation is always initiated with 500 individuals (*NIND*) randomly placed on the landscape. The level of usability of the habitat is always set to 100% (*N_USABILITY*) and the level of usability of the hostile matrix is always set to 0% (*H_USABILITY*). The level of usability of the suitable matrix is a given number between 10% and 90% (*USABILITY*). An individual dwelling in a cell with a usability level that equals 100% will be facing the exact same breeding habitat described in Fahrig’s model. Therefore, the probability of movement / death / reproduction per time-step of this individual is the same, 0.5 / 0.05 / 0.5. In a cell with a usability level that equals 0%, an individual will be facing the exact same matrix described in Fahrig’s model. Therefore, the probability of movement / death / reproduction of this individual will be 1.0 / 0.5 / 0.0 (see Figure 1).

To analyse in which situations the matrix quality could mitigate the negative effect of fragmentation we run multiple simulations with systematic combinations of the parameters related to matrix quality and fragmentation: *FRAG, USABILITY* and *COVERM*. The rest of the parameters related to landscape profiles were held at the default value (see TABLE 1). These values were selected as default for this analysis because they generate landscapes for which the effect of fragmentation is the highest. Therefore, without increasing the matrix quality the effect of fragmentation exists (as depicted in Figure 3) in any of the landscapes we generated.

### Systematic combination of Fragmentation and matrix quality

To systematically analyse the effects of FRAG, *COVERM* and *USABILITY*, we held the proportions of habitat in the default value (10% of the landscape cells). Thus, in all of our simulations, the overall matrix area was 90% of the landscape cells. We ran simulations with 9,000 different landscape profiles: 900 matrix profiles were generated by the systematic combination of *COVERM* (0.00 to 0.99, increment of 0.01) and *USABILITY* (10% to 90%, increment of 10%). For each of these matrix profiles, we generated landscapes with 10 different levels of fragmentation FRAG(0.1 to 1.0, increment of 0.1). Because of the probabilistic nature of the landscape building routine and individual routines, for each landscape profile we ran 15 simulations replicates, resulting in a total of 135,000 simulations in the entire study.

At the end of each simulation, we recorded in which time-step the population went extinct and the landscape profile. Thus, for each landscape profile, there was a mean of 15 populations’ extinction time. If the population was not extinct until the 500th time-step, the survival time recorded was 500.

## Results

We observed that of the 900 combinations of matrix profiles, the fragmentation effect was present in 513 (57%). Therefore, in this analysis, increases in the overall matrix quality could suppress the negative effects of fragmentation in 43% of the situations (Figure 4).

**Figure 4.**
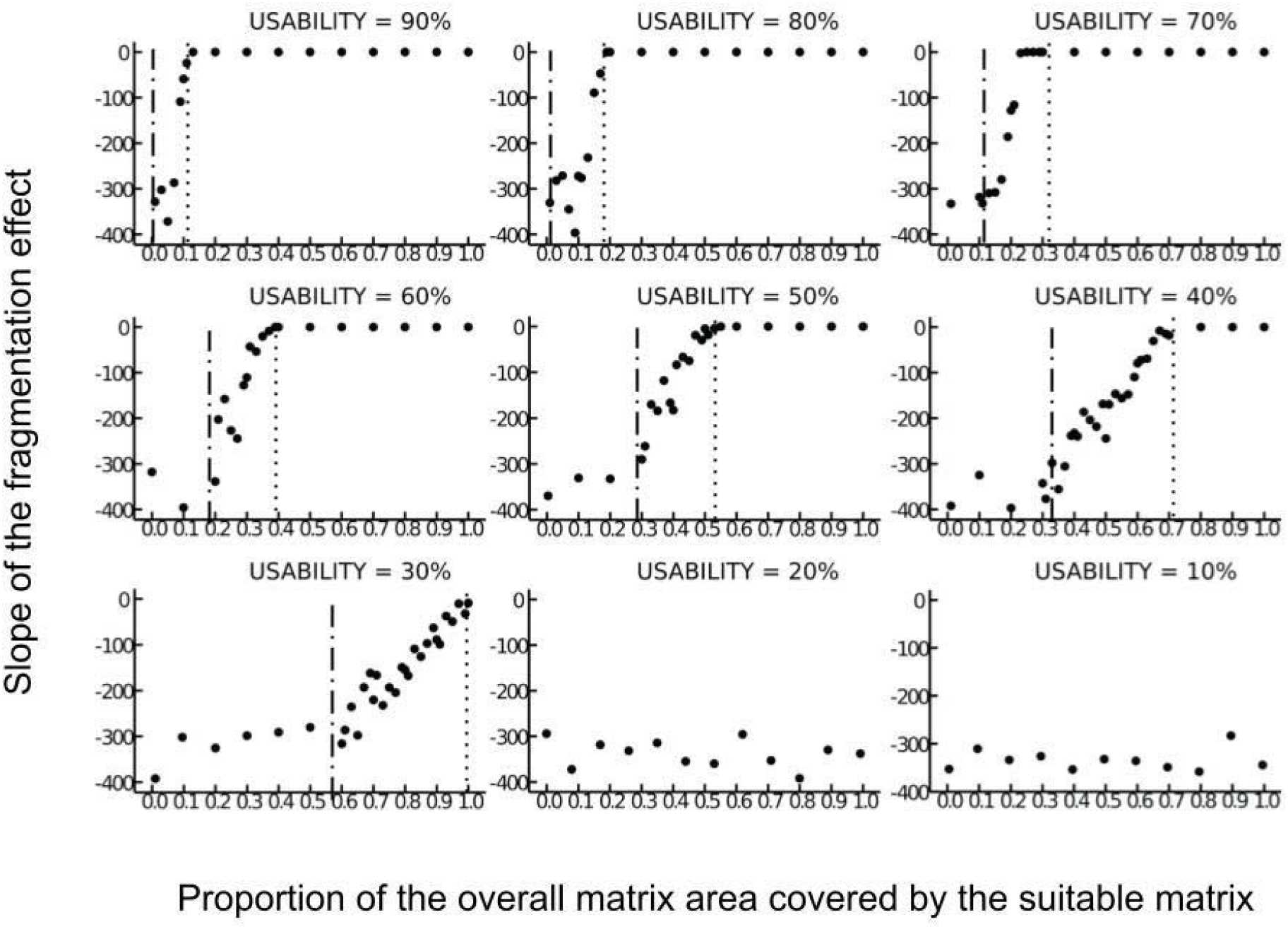
Effect of increasing the quality of the matrix on the fragmentation effect. Each of these graphs shows how much of the overall matrix area needs to be covered by the suitable matrix (as opposed to the hostile matrix) to mitigate the fragmentation effect. The dotted lines show the proportion of suitable matrix needed to completely suppress the fragmentation effect for each level of usability (suppression threshold). The dash-dotted lines show the proportion of the suitable matrix needed to start mitigating some of the fragmentation effects (mitigating threshold). Each point in a graph represents the slope of the fragmentation effect (as depicted in Figure 3). To create each of these graphs, we ran a fragmentation analysis for each value of *COVERM* (represented in the X-axis) resulting in 150 simulations per point. This procedure was conducted for all 9 values of *USABILITY* of the suitable matrix.

The main result of this analysis is that the fragmentation effect can be mitigated and even suppressed by either increasing the proportion of the suitable matrix (*COVERM*) or by increasing the usability of the suitable matrix (*USABILITY*). However these two parameters interact in non-trivial ways, revealing thresholds in which the proportion of suitable matrix begins mitigating the fragmentation effect (mitigating thresholds) and thresholds of proportion above which the fragmentation effect is completely suppressed (suppression thresholds) (see vertical lines in Figure 4). If the proportion of the suitable matrix in the landscape reaches the mitigating threshold it means that the fragmentation effect begins to be reduced at that level. If the proportion of suitable matrix reaches the suppression threshold, it means that the effect of fragmentation is completely suppressed and that the populations always survive for an undetermined amount of time regardless of habitat fragmentation levels. The higher the usability level of the suitable matrix, the lower the mitigating and suppression thresholds. For example, with a usability level of 30% the mitigating threshold is 0.6 and the suppression threshold is 1.0. On the other hand with a usability level of 90%, the mitigating threshold is close to 0.0 and the suppression threshold is 0.12 (see dotted lines in Figure 4).

There are situations in which increases in the proportion of the suitable matrix does not suppress or even mitigate the fragmentation effect For example, the graphs showing levels of usability 20% and 10%, reveals situations in which the similarity of the suitable matrix with the habitat are so low that it cannot suppress the fragmentation effect regardless of its proportion in the landscape and regardless of being more usable than the hostile matrix.

As the level of usability of the suitable matrix increases, the distance between the mitigating and the suppressing thresholds decreases. This implies that at higher levels of usability, a small increase in the proportion of suitable habitat will have a greater impact on survival time than the same increase in proportion for lower levels of usability.

## Discussion

Our results show that when habitat loss is controlled and the level of fragmentation of habitat is determinant of population permanence in the landscape, the existence of patches of suitable matrix can improve populations’ survival time by mitigating and even completely suppressing the negative effect of fragmentation. However, the effectiveness of the suitable matrix in reducing the effects of fragmentation per se depends on their level of usability and their proportion in the landscape. Matrices with higher usability levels require less coverage to entirely suppress the negative effects of fragmentation, while matrices with lower usability levels either require large amounts of coverage for the same result, or simply are not able to suppress fragmentation’s effects regardless of its coverage.

There are two main biological explanations for how matrices can mitigate the effect of fragmentation *per se*. First, as the habitat is broken apart in several patches, suitable matrices may maintain landscape connectivity between these patches by allowing individuals to transit between them (Bélisle, 2005). Therefore, a fragmented landscape with a more suitable matrix can offer more connectivity between patches than a fragmented landscape with only a hostile matrix due to the positive effects of on movement and dispersal between habitat patches (Driscoll et al. 2013; Prevedello et al. 2016). The exchange of individuals among patches reduces the isolation effects, increasing (re)colonisation rates and thus reducing the chances of population extinction (Prevedello et al. 2016; Boesing et al 2018a). Second, organisms can use the matrix to supplement or complement resources that are less available in isolated patches of native habitat (Ewers & Didham, 2006). As a consequence, isolated habitat patches neighbouring suitable matrices can support bigger populations than isolated habitat patches neighbouring hostile matrices (Hatfield et al. 2020). In both cases, if the suitable matrix is able to compensate for the isolation and reduced size of habitat patches caused by fragmentation, the negative effect of fragmentation on the population permanence in the landscape ceases.

The effectiveness of the mechanisms described above is shaped by the usability level of the suitable matrix. The biological explanation behind what we observed is that matrix types structurally more similar to native habitat may offer less movement resistance to the species, increasing functional connectivity in the landscape, and thus being considered of better quality (Prevedello and Vieira, 2010; Estavillo et al., 2013; Boesing et al. 2018b). Matrices more similar to habitat and, therefore, with higher levels of usability can increase species cross-habitat spillover due to reduction in edge contrast with habitat (Rejinfo, 2001), encouraging habitat-dependent species to move between habitat patches (Boesing et al. 2018a). Boeing et al. (2018b) have shown that matrices composed of sun coffee plantations are much better in preventing biodiversity loss and, consequently, delaying biodiversity loss thresholds in neighbouring forests than matrices composed solely of pasture. Similar results have also been found for landscapes with matrices composed of eucalypt plantations but of different ages (Calviño-Cancela et al. 2012; Rocha et al., 2013). According to the authors, the age of the forest reflects its similarity to the forest as it increases the density and diversity of the understory. Their result shows that the older the plantation, the higher the diversity of forest species due to higher connectivity between natural habitat patches. Therefore, eucalypt plantations with different ages may offer different levels of usability. Similar to the results from our models, empirical evidence suggests that matrices structurally more similar to habitat and thus of higher quality (e.g., sun coffee plantations and older eucalypt plantations) can be a useful strategy to maintain populations in fragmented landscapes by reducing edge effects and offering similar resources to the ones in the habitat (e. g. nesting sites, protection from predators, fruits, protection from the sun) (Prevedello and Vieira, 2010; Estavillo et al., 2013; Boesing et al. 2018a;b). Despite patch size and isolation consistently having a stronger effect on biodiversity than matrix type (Prevedello and Vieira, 2010), increasing matrix quality may act as an alternative to mitigate the negative effects of fragmentation. Past and present land conversions have caused a great deal of biodiversity loss and, when restoration is possible (see discussion in Choi, 2007), recovering natural habitats would sometimes require longer than the time needed for some species to be extinguished in the landscape (Dobson et al., 1997). Our study shows that even when habitat configuration properties (patch size and isolation) are controlled (see Figure 2), there are scenarios in which the matrix type can prevent population’s extinction in the landscape.

Our results also establish that there is an interaction between the level of usability of the matrix and the amount of coverage needed to completely suppress the negative effect of fragmentation. The higher the level of usability of the suitable matrix, the less area of this matrix type would be needed to completely suppress the effects of fragmentation on population permanence in the landscape. For example, for forest dependent species, the number of bird populations a patch of forest can maintain is higher than what would be expected from a sun coffee plantation patch of the same size (Boesing et al. 2018b). This is due to the fact that birds are known to partition canopy strata (MacArthur & MacArthur, 1961) and forest patches have more diversified canopy strata than coffee plantations (Boesing et al. 2018b). In other words, to offer the same amount of bird shelter a patch of forest does, a landscape would have to have a much larger area covered by sun coffee plantations. This result highlights the need to consider not only matrix quality but also the amount of it needed in the landscape to truthly address the negative effects of fragmentation. Nevertheless this relationship might not be linear.

Below a certain level of usability, no increase in the proportion of such a matrix type would be enough to suppress or even mitigate the effect of fragmentation. Although there are no empirical studies showing the systematic effect of matrix quality and its proportion in the landscape on population’s survival time, there are empirical patterns that point towards the mechanism revealed in our model. For example, different studies have considered eucalypt and conifer plantations more usable for forest species then other kinds of matrix such as pastures, urban areas, and arable land (Barlow et al., 2007; Fonseca et al., 2009; Calviño-Cancela et al,. 2012; Rocha et al., 2013). Nevertheless, in all these studies, there were always species or functional groups that were never found inside these matrices. This implies that, however structurally similar such plantations are to forests, they are just not similar enough for some species to carry out essential life processes. In this scenario, what our model predicts is that, no matter how many patches of, for example, pasture areas are converted to eucalypt plantations in this landscape, for some species, this change will not reduce the risk of extinction caused by habitat fragmentation. This result points towards the idea that heterogeneous matrices can be the best way to maintain biodiversity because they provide a much higher variety of resources (Chetcuti et al., 2021).

Therefore, considering matrix quantity along with quality has relevant implications for landscape management. In Brazil, matrices composed of monocultures (i. e. pasture and soy plantations) have shown to be very hostile causing great loss to forest biodiversity even with the preservation of 20% of native forest demanded by law (Rigueira et al., 2013). With the assumption that only a small portion of the native habitat will remain intact, it is especially important to understand how matrix quality can help maintain biodiversity. Although large-scale, higher-quality matrix conversion may not be realistic at the present (Boesing et al. 2018b), it is important to have in mind that there are experiences with agricultural initiatives different from monocultures, such as agroforests, that have improved biodiversity (Boshier 2004; Hagar et al. 2019). Therefore, although there is evidence that matrix heterogeneity improves biodiversity (Chetcuti et al., 2021), our model shows specifically what to expect from combinations of quality and quantity of elements of this heterogeneity. Additionally, these differences in outputs from combinations of matrix quality and quantity can help explain some of the reported variability in the effects of fragmentation per se on biodiversity (Fahrigh et al. 2019).

Future research could look into the creation of a land-use association index that would estimate species-specific or functional group-specific levels of usability for different land uses, while accounting for the amount of it that is needed in the landscape. With such an index, it would be possible, for example, to parameterize models like ours to pinpoint the exact values of similarity to habitat and proportion needed to completely suppress negative effects of fragmentation and habitat loss. These indexes might not be so far in the future as some might fear. Potentially useful data such as species occurrence and abundance and habitat dependency are becoming more and more available in the literature (Chetcuti et al. 2019) and there are indications that structural complexity can be a good proxy of similarity between habitat and matrix (Prevedello and Vieira, 2010; Calviño-Cancela et al., 2012; Rocha et al. 2013; Boeing et al., 2019). Nevertheless, having such an index of similarity between matrix types and native habitats would still take a lot of effort and knowledge. Identifying what kind of matrix and at which proportion it is more efficient in maintaining natural processes is not an easy task mainly because “usability” can be group specific. For example, eucalypt plantations may be more similar to tropical forests in structural complexity than sun coffee plantations (Calviño-Cancela et al., 2012) and therefore offer more nesting and protection sites. However, coffee plantations may temporarily offer higher availability of carbohydrates in berries and nectar of flowers, which may be more attractive for species while foraging outside the habitat (Boezing et al. 2018b). Therefore other biological aspects aside from structural similarity, such as functional traits of focus species, should also be taken into account.

In synthesis, our results show that not only matrix matters but the quality and proportion of it matters too. The more similar the matrix is to the habitat the more it matters, and the less coverage of it is needed to preclude negative effects of habitat fragmentation. Although still suitable, some matrices offer so little usability that no amount of such matrices is capable of suppressing or even mitigating fragmentation effects on population persistence. Our work not only reinforces the need to account for matrix quality when evaluating fragmentation effects in real-world, complex landscapes, but also highlights the essential role the amount of suitable matrices plays in determining the magnitude of the negative effects of fragmentation. An adequate index of similarity between matrix and natural habitat would greatly improve our efforts to reduce negative effects of habitat fragmentation and loss via matrix management. Finally, future studies should also focus on how combinations of matrices with different qualities but also quantities shape fragmentation’s effects on population persistence, and thus help explain the reported variability in the effects of fragmentation per se on biodiversity.

## Acknowledgements

We thank L Fahrig for her kind help with our doubts about her model and for giving us the permission to use one of the pictures of her work. B Travassos-Britto was supported by Coordenação de Aperfeiçoamento de Pessoal de Nível Superior (CAPES) (grant number 88882.315632/2019-01). C Hohlenwerger was also supported by CAPES (88882.327885/2019–01 and 88887.309513/2018–00). PBL Rocha was supported by the Fundação de Amparo à Pesquisa do Estado da Bahia (FAPESB). PBL Rocha and JGV Miranda were supported by Conselho Nacional de Desenvolvimento Científico e Tecnológico (CNPq) during this project.

