## Supporting information S1 for "Quantity and quality of suitable matrices matter in reducing the negative effect of fragmentation on populations extinction risk"

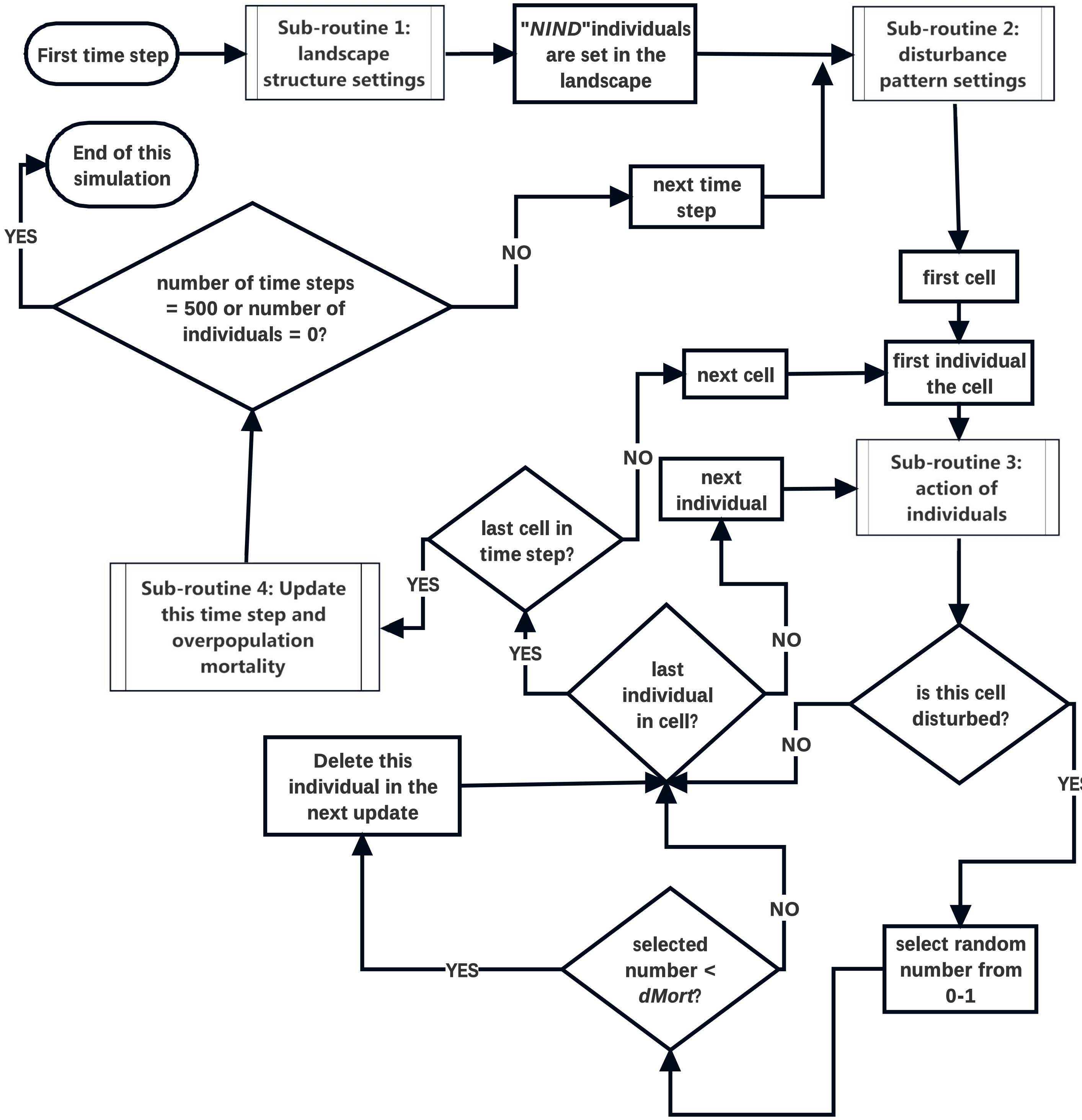


Main routine. Algorithm to be executed by the software to run a single simulation.

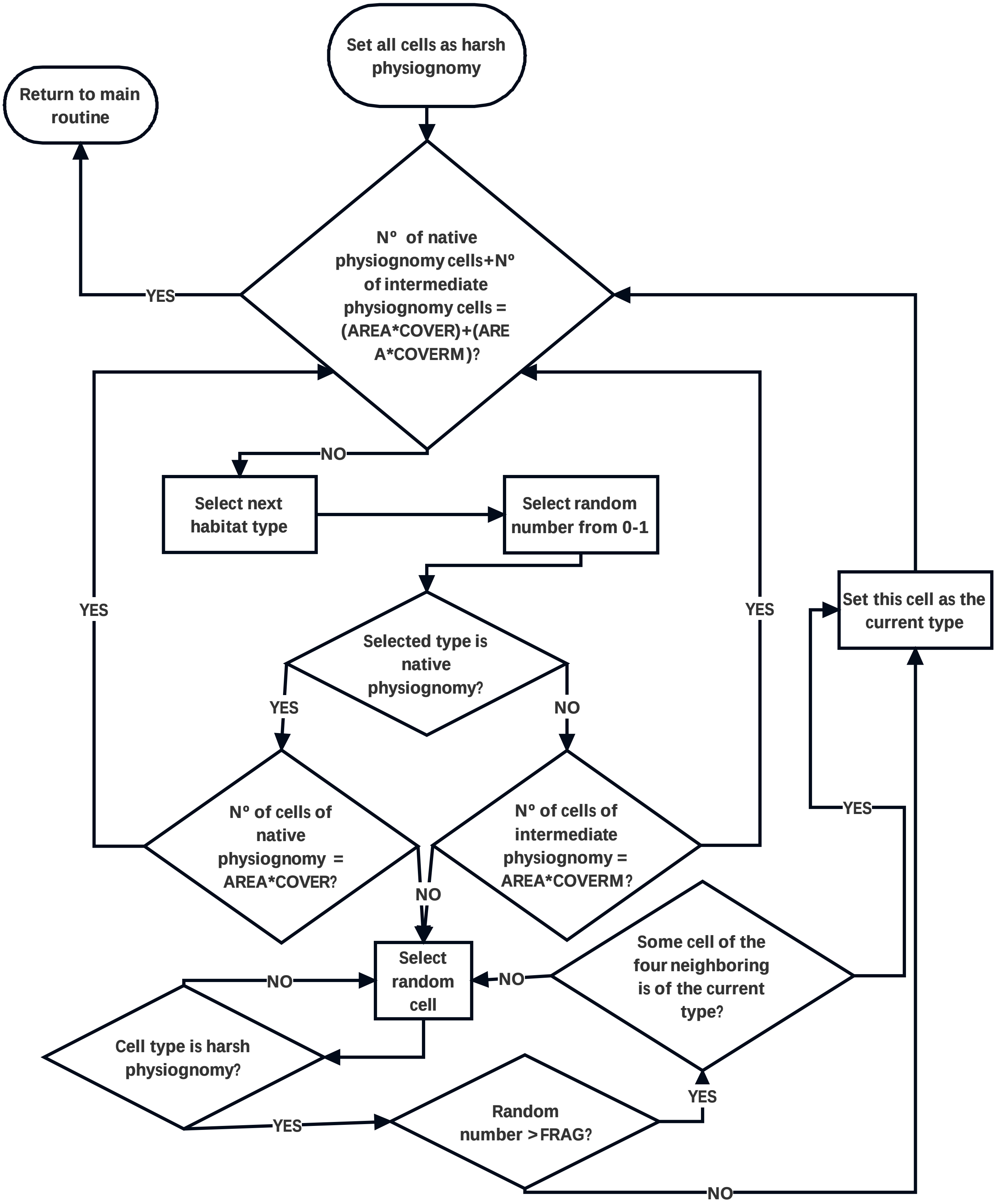


**Sub-routine 1. Algorithm setting properties of cells and consequentially the landscape design.**

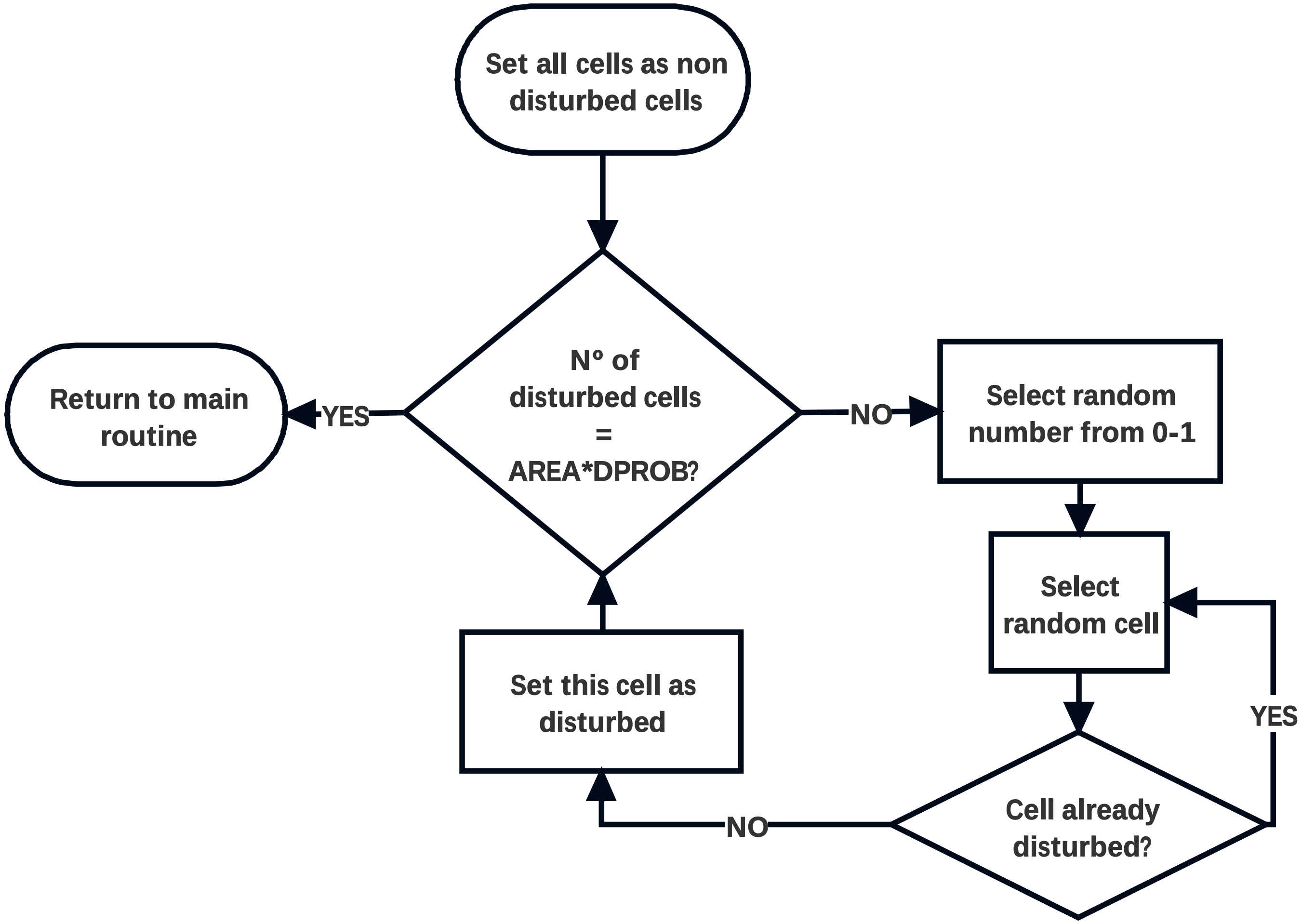


**Sub-routine 2. Algorithm to set the disturbance pattern in each landscape.**

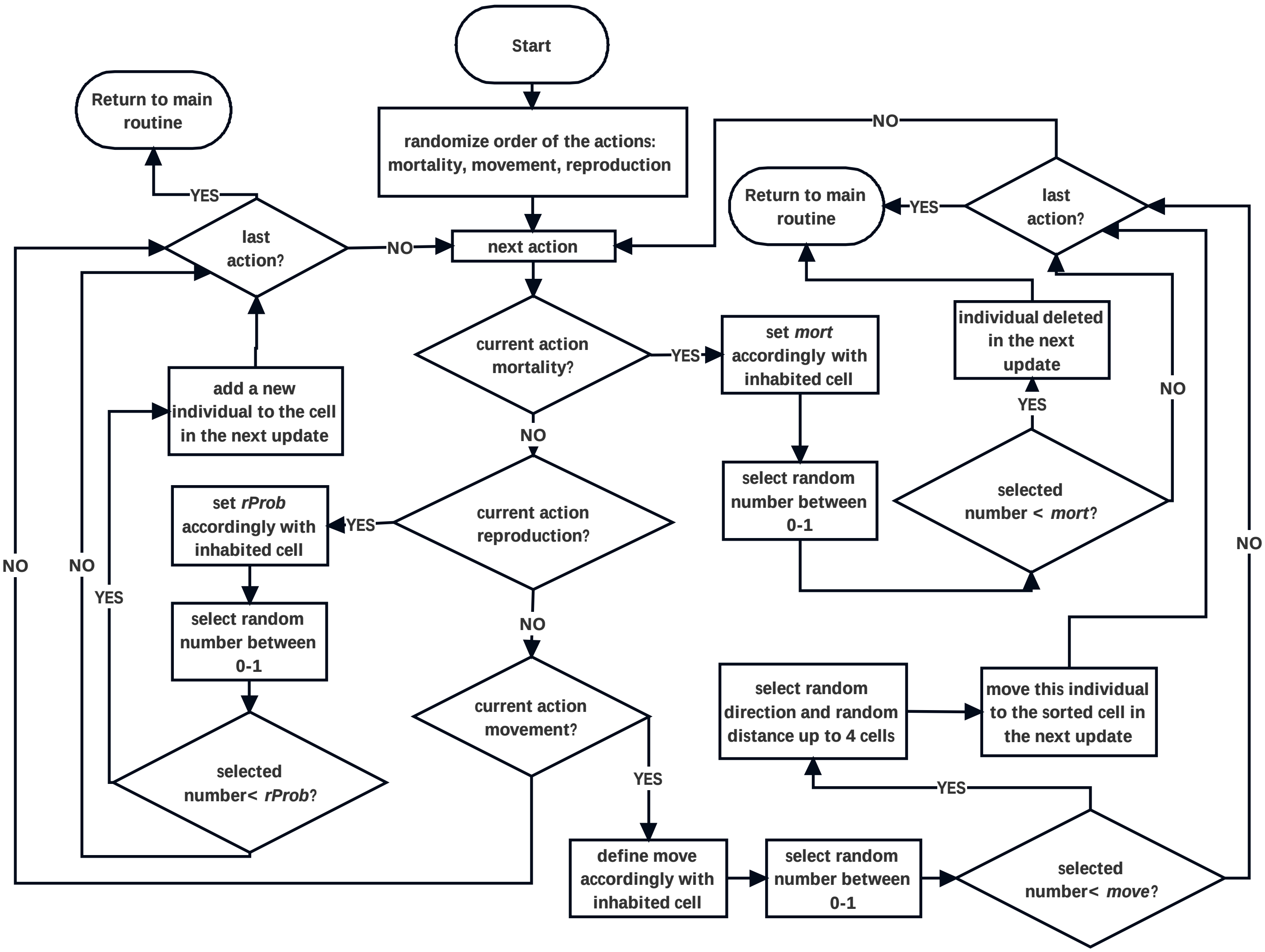


**Sub-routine 3. Algorithm followed by each individual in each time step.**

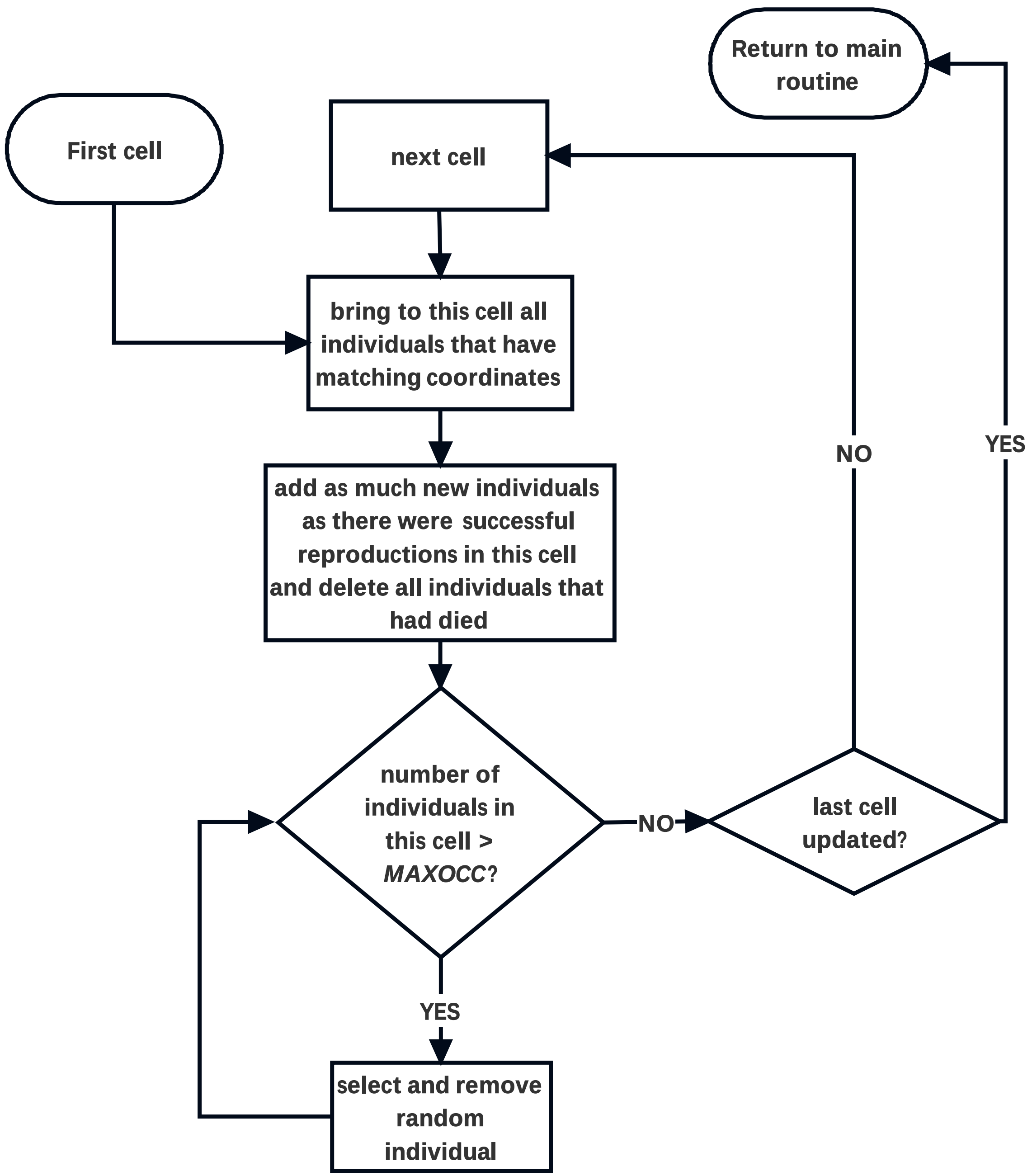


**Sub-routine 4. Algorithm executed by each cell to update death, birth and movement, and death by overpopulation in the cells.**
