## Supporting information S2 for "Quantity and quality of suitable matrices matter in reducing the negative effect of fragmentation on populations extinction risk"

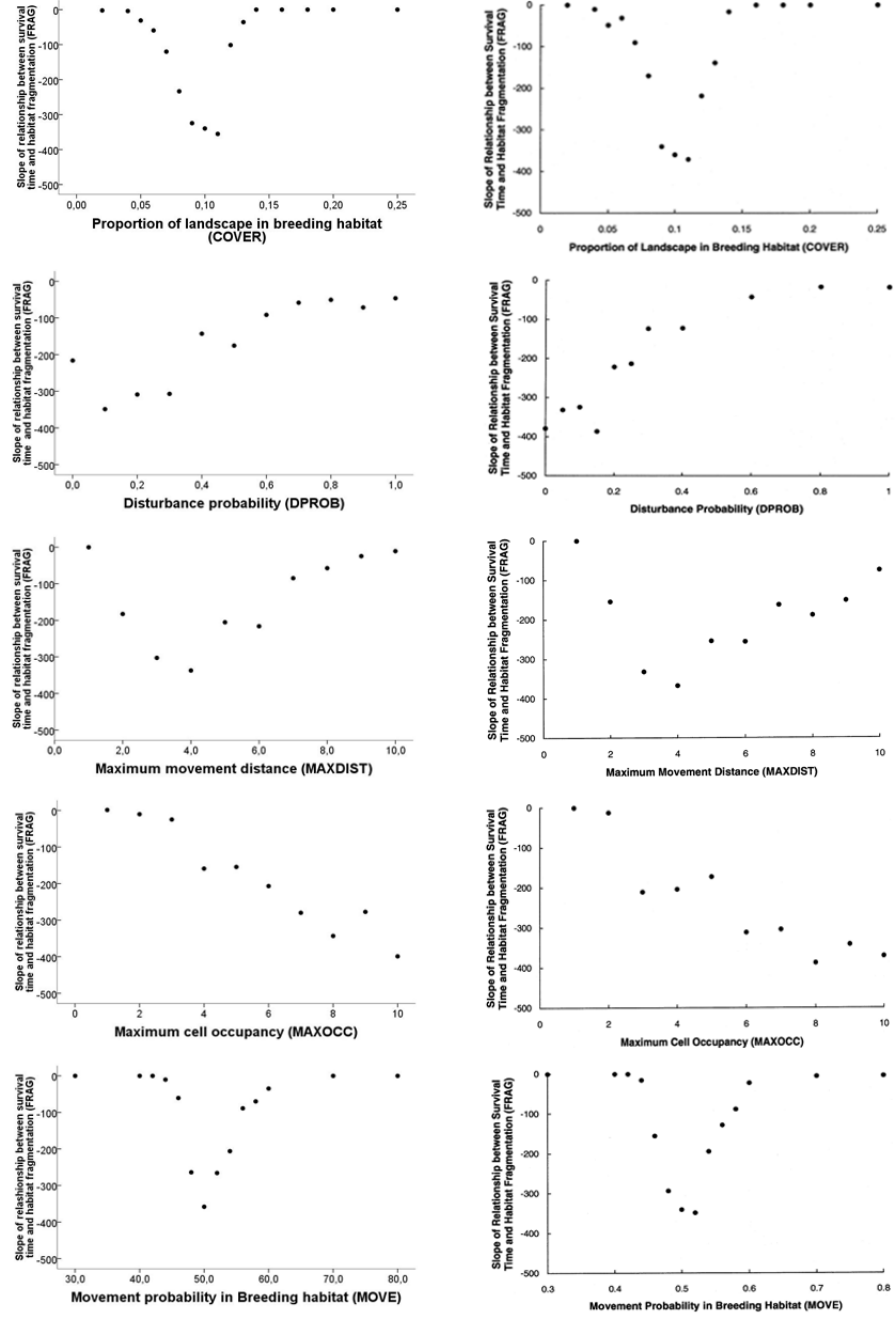


Comparisson between results of Fahrig's model (1998) (rigth) and our model (left) with relation to variations in specific parameters (Fahrig, 1998 – Figure 8, 9, 557 12, 14, 15).
